# Humans Optimally Integrate Cutaneous and Proprioceptive Cues In Haptic Size Perception

**DOI:** 10.1101/2025.06.24.661403

**Authors:** Keon S. Allen, Daniel Goldreich

## Abstract

Sensory perception often relies on the brain’s integration of multiple noisy inputs (cues), a process known as cue combination. Cue combination within the sense of touch has been understudied. Here, we investigated whether humans optimally combine haptic cutaneous and hand configuration cues when discerning the size (e.g., diameter) of a disk held edge-on between the thumb and index fingers. When these two fingers span the diameter of a disk to contact its perimeter, a hand configuration cue (relating to the perceived distance between the fingers) provides information about the disk’s size. Less obviously, cutaneous cues to disk size may be provided simultaneously from the indentation of the skin caused by the curvature of the disk (smaller disks cause greater indentation). It is unknown whether humans make use of all these cues when perceiving the size of the held object, and if so, whether they integrate the cues optimally. We considered three hypotheses for how humans might use these cues: they might rely solely on the least noisy cue (Winner-Take-All Model, WTA), combine cues based on a simple arithmetic average (Average-Measurement Model, AVG), or combine cues via an optimal weighted average (Optimally-Weighted Model, OPT). In three experiments involving 34 participants, we measured the reliabilities of these cues and compared participant performance to the predictions of the three models. Each experiment tested participants using a two-interval forced-choice (2IFC) paradigm with 3D printed disk stimuli. On each trial, under occluded vision, participants felt two disks sequentially and responded which felt larger. Participants were tested with each finger’s cutaneous cue alone, the configuration cue alone, and all three cues together. In two experiments, the disks presented were circular. In a third experiment, unknown to participants, some of the presented disks were oval-like cue-conflict stimuli. The improvement of accuracy observed in multi-cue conditions over single-cue conditions, and the Point of Subjective Equality (PSE) shifts observed in cue-conflict conditions, were consistent with optimal cue combination. We conclude that humans are capable of combining haptic cutaneous and configuration cues optimally to judge the sizes of held objects.

## 1 Introduction

Touch, the first sense to develop in humans [1, 2], facilitates our interactions with the world and is critical for manipulating, interacting with, and experiencing our environment. Anything felt must be translated from interactions with the skin to patterns of neural impulses that are subsequently decoded by the brain. As the skin is a uniquely movable and deformable organ, the brain presumably incorporates the configuration of the limbs and digits in space, in addition to various cutaneous inputs, in order to interpret the shape of a held object. How does the brain combine these distinct haptic cues in order to achieve a final percept?

Previous cue combination research has generally focused on the integration of cues arising from two or more senses, or on the integration of two or more cues within non-haptic senses. Alais and Burr [3] found that, when localizing a source that emitted a beep and a flash, participants performed more accurately with both cues than with either alone. Hillis et al. [4] similarly found that participants more accurately estimated stimulus depth when using two visual cues, texture and binocular disparity, rather than either alone. Crucially, many studies in these domains have found that, during the perceptual process, the brain weights cues according to their reliability [4–11], a hallmark of optimal cue combination [12, Chapter 5].

The integration of cues within the sense of touch has been relatively understudied, and it is currently unclear whether human performance during haptic exploration conforms to an optimal cue combination strategy [13–15]. Here, we investigated haptic cues relating to the sizes of different coin-like disks held edge-on between the thumb and index fingers. When feeling for a coin without looking, humans can discern different denominations by their sizes. Clearly, the size of the coin relates to where the fingers are in space through a proprioceptive signal. In order to hold a larger coin, the fingers must be spread farther apart; thus, disk size is presumably signaled at least in part by this hand *Configuration Cue (Config)*. Less obviously, a coin held on its perimeter between the index finger and thumb exerts pressure that indents the skin of each finger. Given a particular contact force, the indentation profile relates to the size of the coin through its curvature. Smaller coins have greater radii of curvature and produce correspondingly greater skin deformation, which is signalled by the firing rates of slowly adapting tactile afferent axons [16]. We refer to these cutaneous size cues from the thumb and index finger as *Cutaneous Cue 1 (Cut*.*1)* and *Cutaneous Cue 2 (Cut*.*2)*, respectively. Do humans combine *Cut*.*1* and *Cut*.*2* with *Config* in order to perceive disk size? If these cues differ in their reliabilities, do humans weight them optimally during the perceptual process?

We investigated three hypotheses for how humans might utilize these haptic cues to perceive the sizes of disk-shaped objects. Perhaps the brain bases its perception exclusively on the most reliable cue and ignores the others. We refer to this possibility as the *WTA* model. Alternatively, perhaps the brain does integrate the three cues but without taking into account their differing reliabilities, instead basing perception on the simple arithmetic average of the cues. We refer to this possibility as the *AVG* model. Finally, perhaps the brain indeed weights each cue according to its reliability, basing perception on an optimal weighted average of the cues. We refer to this possibility as the *OPT* model.

Mathematically, the models’ distinct perceptual strategies can be represented by the following percept formulae. In these formulae, we denote the stimulus, disk size (mm radius), by *s*, the standard deviation of a cue across repeated trials by *σ*_*Cue*_, the most reliable cue by *Cue\**, and the percept (perceived size of the disk size) by *ŝ*. For future reference, we define the *reliability* of a cue as equal to the cue’s inverse variance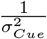.

WTA:

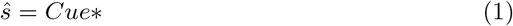

AVG:

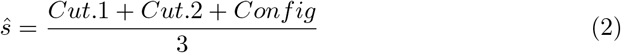

OPT

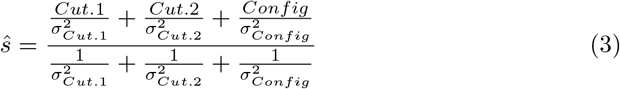

With reference to the above percept formulae, note that, when one of the cues is exceptionally reliable (i.e., its sigma is much smaller than the others), the *OPT* and *WTA* models make similar predictions. As the sigma of a cue approaches zero, that cue is increasingly relied upon by an optimal observer, and consequently the *OPT* model reduces to the *WTA* model. When the sigmas of the three cues are similar, the *OPT* and *AVG* models give similar predictions, and indeed if the three cues are equally reliable, the *OPT* model reduces to the *AVG* model. When the cue reliabilities vary, the three models makes different predictions. Thus, it is possible to distinguish among the three models, depending on the relative sigmas of the cues.

## 2 Methods

### 2.1 Simulations and Predictions

#### 2.1.1 Model Predictions

We used Monte Carlo simulations to explore the predictions from each model. In a simulated 2-interval forced-choice (2IFC) task, a model participant (simulant) was presented sequentially with two disks of differing size and answered which felt larger. On each trial, one of the disks was a reference disk, while the other was a comparison disk. The size of the comparison disk varied across trials, with a radius from −5mm to +5mm, in increments of 0.1mm, relative to the fixed reference disk radius. In each interval, when a disk was presented, the model’s ‘nervous system’ drew a sensory measurement (cue) from each of three Gaussian distributions centred on the disk radius, with a sigma corresponding to the reliability of the corresponding cue. Each model then processed the three cues according to its particular percept formula (above). On every trial, the simulant reported which of the two disks it perceived to be larger. We simulated 1000 trials at each comparison level. The proportion of times the comparison disk was judged larger than the reference disk is plotted to create a psychometric function for each model (Figure 1).

**Fig. 1.**
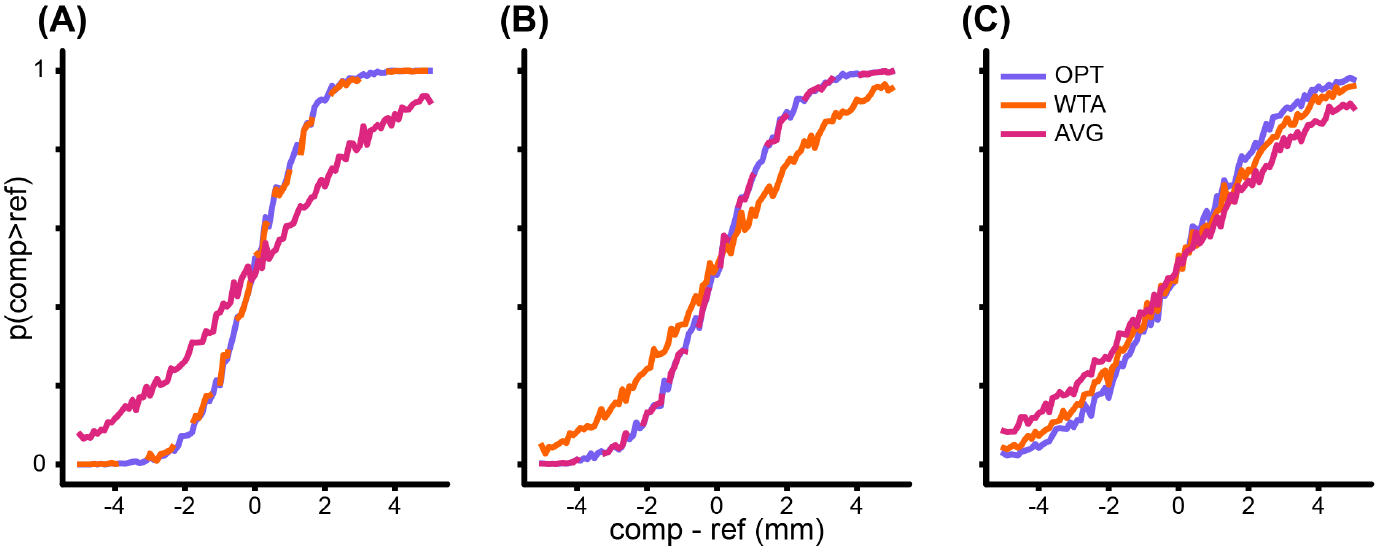
Monte Carlo simulations, from a single simulant, of model predictions under different relative sigma values. For each of the three models, in each panel, there were 1000 simulated trials at each 0.1mm increment from −5mm to +5mm. **(A)** When one cue has a much smaller sigma than the others (ie. 8mm, 8mm, 2mm), *OPT* and *WTA* models make similar predictions. **(B)** When the cues have similar sigmas (ie. 4mm, 4mm, 4mm), *OPT* and *AVG* models make similar predictions. **(C)** When the relative sigmas vary (ie. 7mm, 6mm, 3mm), the models makes different predictions.

Importantly, note that the three models make distinct predictions regarding the variability of the percept (i.e., the standard deviation of the percept across trials), which is reflected in the slope of the corresponding psychometric functions. From the models’ percept equations, above, it is straightforward to derive the standard deviation of the percept across trials when the same disk is felt repeatedly:

*WTA:*

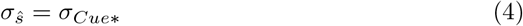

*AVG:*

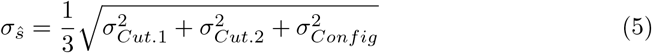

*OPT:*

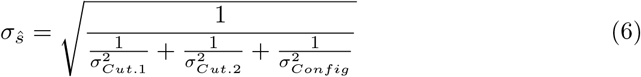

Note that the *OPT* model percept is less variable than that of the other two models.

### 2.1.2 Experiment Simulations

In order to validate our categorization methods and to assess their accuracy, we simulated a realistic Method of Constant Stimuli (MCS) two-interval forced-choice (2-IFC) experiment with thousands of simulants who combined cues according to each of the three models. We considered three different populations of simulants defined by the relative sigmas of the sensory cues. Within each population, the sigma for each cue was normally distributed (with mean *M*_*Cue*_ and standard deviation *S*_*Cue*_); to create a simulant, the sigma for each cue was sampled from the corresponding normal distribution.

The three population were as follows:

Population 1: the sigma of one cue was exceptional (*M*_*Cut*.1_ = *M*_*Cut*.2_ = 8*mm, S*_*Cut*.1_ = *S*_*Cut*.2_ = 2*mm*; *M*_*Config*_ = 2*mm, S*_*Config*_ = 1*mm*).

Population 2: the sigmas of the three cues were similar (*M*_*Cut*.1_ = *M*_*Cut*.2_ = *M*_*Config*_ = 4*mm*; *S*_*Cut*.1_ = *S*_*Cut*.2_ = *S*_*Config*_ = 0.1*mm*).

Population 3: the sigmas of the three cues differed (*M*_*Cut*.1_ = 7*mm, S*_*Cut*.1_ = 2*mm*; *M*_*Cut*.2_ = 6*mm, S*_*Cut*.2_ = 3*mm*; *M*_*Config*_ = 3*mm, S*_*Config*_ = 1*mm*).

The simulated experiment was split into four conditions. In each condition, there were 8 comparison levels with 20 trials per comparison level. In the first three conditions, the simulants used each individual cue alone; in the fourth condition, the simulants performed the same task using all three cues together.

On a given trial, the simulant compared a reference stimulus with a comparison stimulus. In the Combined Condition, for example, the measurement distribution for each cue was a Gaussian centred on the true radius of the piece, with a standard deviation equal to the sigma a given simulant had for that cue. In each interval of a trial, the three individual cues were combined according to one of the model equations. The simulant determined which of the values in each interval was larger in order to make its ‘response’ and end a simulated trial. The proportion of times the comparison was perceived greater than the reference formed a simulant’s psychometric function, which we estimated as a cumulative normal distributions of the stimulus level, Δ (Eq. 7).

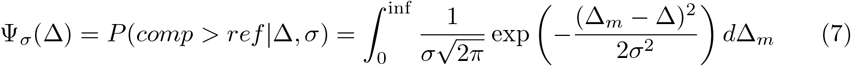

Here, Δ_*m*_ refers to the simulant’s internal measurement of the size difference between the two disks, which results from a random sample drawn from a Gaussian distribution centered on the actual size difference, Δ.

The performance data (D) for each simulant were analyzed as for a human participant. For a given simulant, we used their performance in the three individual-cue conditions to calculate predictions for their performance in the Combined Condition according to each of the three models. Specifically, for each single-cue condition, for each simulant, we first calculated the likelihood:

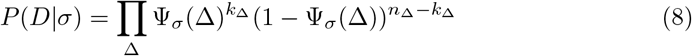

Where *k*_Δ_ is the number of correct trials and *n*_Δ_ is the total number of trials at Δ. We then calculated the posterior *PDF* over *σ*_*cue*_ via Bayes’ formula using a uniform prior *PDF* over sigma:

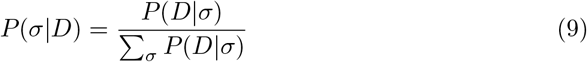

By integrating over the *σ*_*cue*_ *PDF* for each of the individual-cue conditions, we calculated the *σ*_*cue*_ *CDFs*. By uniformly sampling from the ordinate of the *σ*_*cue*_ *CDFs* and interpolating to the abscissa, we sampled sigma values from the *σ*_*cue*_ *PDFs*. This sampling procedure was preferable to using the mode of the *σ*_*cue*_ *PDF* as a point estimate because the spread of samples reflected the uncertainty of the estimate. A single draw from each of the three *σ*_*cue*_ *PDFs*, yielded a triplet that served as input values for each of the three model equations (Equations 4, 5, and 6, above). Sampling a thousand triplets for each participant, we calculated a distribution of Combined Condition sigmas (*σ*_*ŝ*_) for each of the three models, for every simulant. We computed the probability of the simulant’s performance data given each of these 1,000 model-predicted *ŝ*. The average of these 1,000 probabilities implements the marginal likelihood formula for the model, for each participant (Eq. 10).

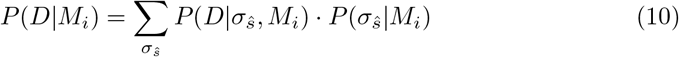

We categorized the simulant as using the model that had the greatest marginal likelihood. The results from this simulated experiment run on 1000 simulants per model are summarized in Table 1.

**Table 1.** Simulant Classification in a Non-Conflict Experiment.

|  | OPT Simulant | WTA Simulant | AVG Simulant |
| --- | --- | --- | --- |
| Classified as OPT | **677** | 516 | 98 |
| Classified as WTA | 315 | **445** | 250 |
| Classified as AVG | 8 | 39 | **652** |

We additionally simulated a cue conflict experiment, in which the reference stimulus in the Combined Condition was no longer a circular disk but rather an oval-like object. The conflict object presented a *Config* cue corresponding to a disk radius that was either larger (positive conflict) or smaller (negative conflict) than the disk radius used to produce the cutaneous cues, *Cut*.*1* and *Cut*.*2*. In this case, the models make distinct predictions about the mean percept *ŝ* (see Equations 1, 2, and 3, above) as well as the *σ*_*ŝ*_ from each triplet. Once again, we computed the probability of the simulant’s performance data given each model by averaging the 1,000 probabilities of the data given each model-predicted (*ŝ, σ*_*ŝ*_). This implements the marginal likelihood calculation for the model 11:

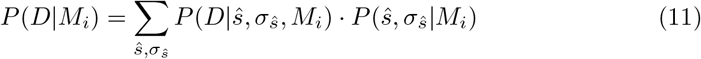

The simulation results run on 1000 simulants per model are summarized in Table 2. Figure 2 shows example data from a single simulant in the simulated cue conflict experiment.

**Table 2.** Simulant Classification in a Cue Conflict Experiment.

|  | OPT Simulant | WTA Simulant | AVG Simulant |
| --- | --- | --- | --- |
| Classified as OPT | **870** | 157 | 10 |
| Classified as WTA | 129 | **843** | 1 |
| Classified as AVG | 1 | 0 | **989** |

**Fig. 2.**
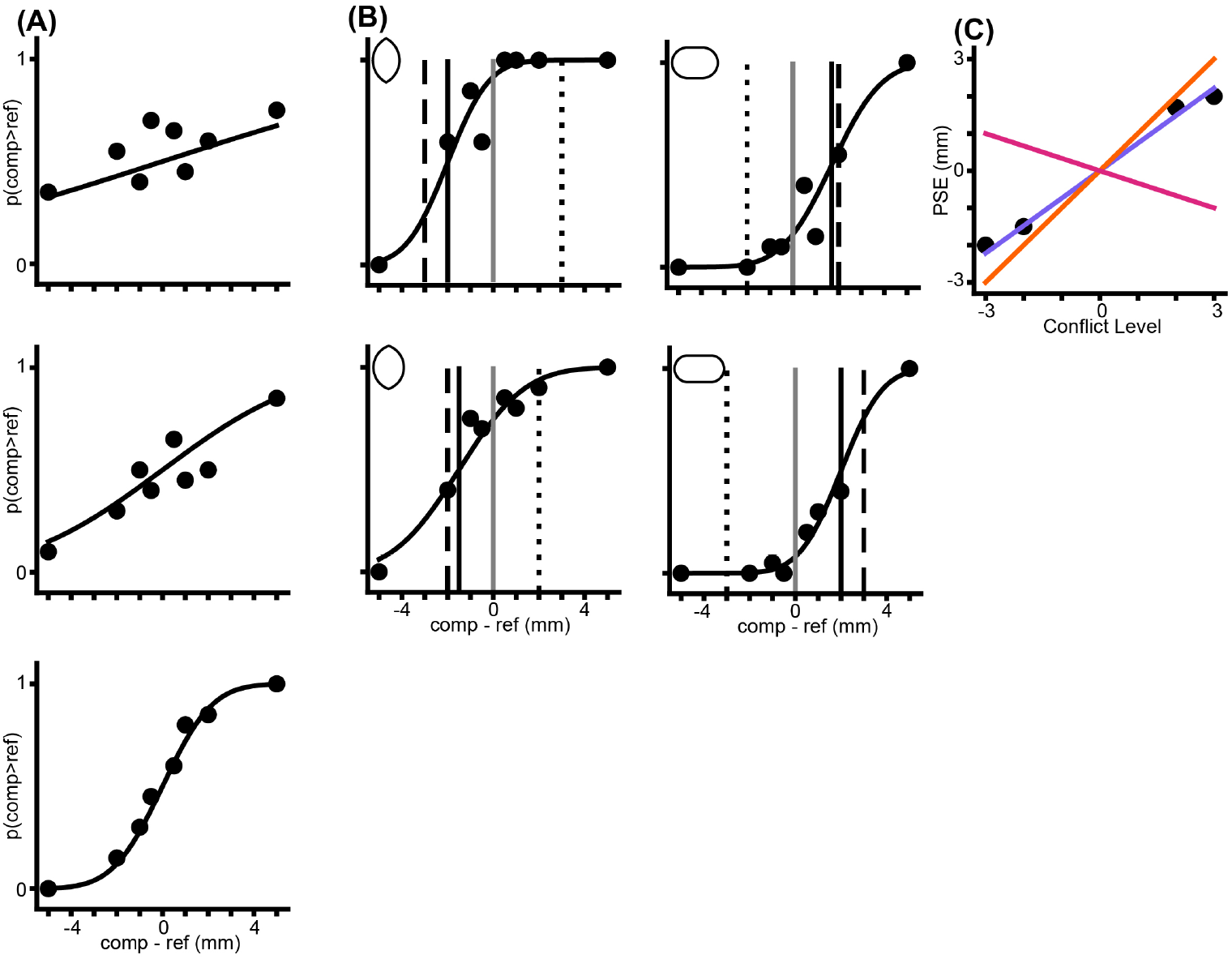
Example of cue-conflict performance from a simulated Bayesian optimal observer with varied sigma. **(A)** Single-cue psychometric functions. From the top: Cut.1, Cut.2, Config. **(B)** Performance under the 4 cue conflict conditions (top left: −3, bottom left: −2, top right: +2, bottom right: +3). Black vertical line: *PSE*. Grey line: zero. Dashed line: performance that would result from exclusive reliance on the Config. cue. Dotted line: performance that would result from exclusive reliance on the two Cut. cues. **(C)** Plot of *PSE* vs. conflict condition, from (B), along with the predictions of the models, based on the estimated single-cue sigmas from (A). Colors as in Fig. 1.

### 2.2 Participants

This study was approved by the McMaster Research Ethics Board. Participants were 34 McMaster University undergraduate students. Persons with any one or more of the following conditions did not participate, as these conditions are known to adversely affect tactile acuity or the ability to perform tactile tasks: diabetes, nervous system disorder or injury (tremor, epilepsy, multiple sclerosis, stroke, etc.), learning disability, dyslexia, attention deficit disorder, cognitive impairment, carpal tunnel syndrome, arthritis of the hands, and hyperhidrosis. Twelve participants took part in Experiment 1 (3 women, 9 men; mean age 19.3 years). Eleven participants took part in Experiment 2 (1 woman, 10 men; mean age 18.8 years). Eleven participants took part in Experiment 3 (2 women, 9 men; mean age 19.1 years). Participants gave signed informed consent and received monetary compensation or course credit for their time.

### 2.3 Materials

We designed all stimuli in OpenSCAD, a 3D script-based modeller. The stimuli were flat coin-shaped disks of 0.54mm thickness, 3D printed with PLA plastic using the Ultimaker 2 Go 3D printer. Disks in Experiment 1 were circular with the following radii in millimetres: 10, 13, 14, 14.5, 15, 15.5, 16, 17, and 20. In experiments 2 and 3, the disk sizes were reduced, as we hypothesized smaller disks would lead to more reliable cutaneous cues [16] while leaving the configuration cue relatively unaffected. Disks in Experiment 2 were circular with the following radii in millimetres: 5, 8, 9, 9.5, 10, 10.5, 11, 12, and 15. As in Experiment 2, comparison disks in Experiment 3 were circular with the following radii in millimetres: 5, 8, 9, 9.5, 10.5, 11, 12, and 15. Experiment 3 also made use of a set of conflict disks with altered shape. In each experiment, disks were fastened to a wheel that rotated to present disks of varying radii for a given trial. This wheel was controlled by an ISM-7411 NEMA 23 National Instruments Integrated Stepper motor that interfaced with the computer via an ethernet cable. The computer ran a custom LabVIEW experiment program that used the LabVIEW SoftMotion module to implement stepper motor commands. Plastic thimbles, used in *Config* trials (see below), were 0.5mm in thickness, designed and 3D printed using the above resources, in a range of sizes such that each participant found two thimbles that fit comfortably over their index finger and thumb. A 3D printed hand rest allowed the thumb and index finger to protrude forward in horizontal alignment while the remainder of the hand was held comfortably stable. The hand rest sat upon an adjustable laboratory scissor jack which acted as a rest and guide for the hand. The jack was affixed to a Heavy Duty 20-inch linear bearing slide rail (Firgelli Automations Canada). The rail allowed participants to slide their hands to the appropriate disk as their arms extended. An FE7B series photoelectric retroreflective infrared light sensor (Honeywell) sent a voltage change to a USB 6210 data acquisition device (National Instruments) when its beam was broken or unbroken by participant movement during trial presentations. Thus, the computer was aware of hand positioning relative to the wheel. The disks, wheel, motor, and other equipment were hidden from participant view behind the white foam core box shown in Figure 3B. Participant responses were recorded with a wireless Bluetooth clicker that interfaced with the experiment program. The experiment program and Bayesian analyses were coded in National Instruments LabVIEW 2018 on PC.

**Fig. 3.**
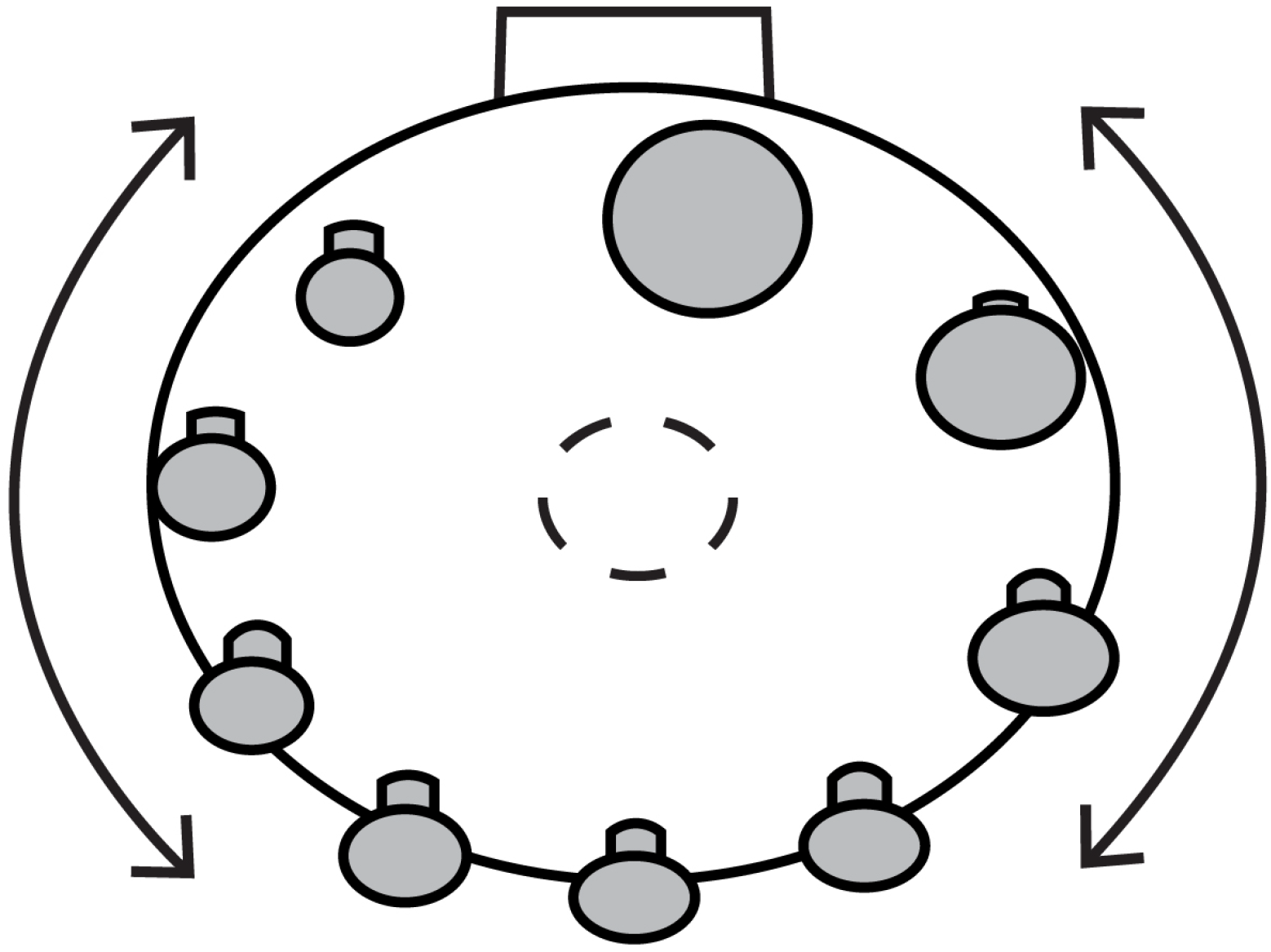
Schematic above-frontal view of the rotating wheel with disk stimuli attached. The wheel is controlled from behind via a stepper motor (not shown). The dashed circle represents the stepper motor axle.

### 2.4 Procedure

Each experiment used a *2IFC* paradigm in order to test participants’ ability to discern which of two disks, felt sequentially, had a larger diameter. The right hand was oriented such that the index finger and thumb were in the horizontal plane. The participants flexed their index finger and/or thumb at the metacarpophalangeal joint in order to touch a disk. On a given trial, a beep signaled the participant to slide their right arm forward to feel the presented target disk. Arm movement was signalled to the computer program by the obstruction of the infrared beam. A second beep, occurring 1000 ms after beam break, signaled the participant to slide their hand back to the initial position. The computer then rotated the stepper motor in order to select a different target disk for the participant to feel. A third beep then signaled the participant to again slide their arm forward and feel this second disk. A final, higher-pitched beep, occurring 1000 ms after beam break, signaled the participant to slide their hand back to the initial rest position. The participant then responded which disk felt larger, by using the Bluetooth clicker in their left hand.

On each trial, one of the two presented disks was a reference disk (presented in every trial) and the other was a comparison disk. The order of presentation of the reference and comparison disks was randomized. Participants completed up to thirty practice trials with feedback before starting each condition.

We implemented controls for timing and distance cues. In order to present a particular target disk, the computer first rotated the wheel to a random disk position, preventing the participant from using timing cues to inform their response. During single-digit experiment blocks, the wheel was calibrated to rotate such that the outer edge of each disk was aligned to be the same distance from the participant’s digit. Thus, the participant could not benefit from proprioceptive or timing cues associated with the flexion of the finger towards the disk. During multi-digit blocks, the wheel was calibrated to keep all presented disks centred at the same position.

Participants in Experiments 1 and 2 were tested in four different conditions measuring performance when they had access to the cues alone and altogether. The four conditions were as follows: the *Cut*.*1* condition, in which participants had access to the cutaneous cue on the thumb; the *Cut*.*2* condition, in which participants had access to the cutaneous cue on the index finger; the *Config* condition, in which participants had access to the hand configuration cue; and the *Combined* condition, in which participants had access to all cues together (Figure 4). In *Cut*.*1* and *Cut*.*2*, participants used only the thumb or index finger, respectively, of their right hand to feel the lateral edge perpendicular to the face of the disk. The disks were left-aligned for the thumb and right-aligned for the index such that the time and distance required by the participant to contact the disk by flexion of the finger, when the hand was in place, did not vary systematically with disk size. In the *Config* condition, participants wore hard plastic thimbles on their index finger and thumb. Thus, as they felt each disk they had access to the *Config* cue but, because the disk was blocked by the thimbles from indenting their skin, they did not have access to either *Cut*.*1* or *Cut*.*2*. In the *Combined Condition* condition, participants did the same task with thimbles removed, having access to both cutaneous and hand configuration cues to disk size.

**Fig. 4.**
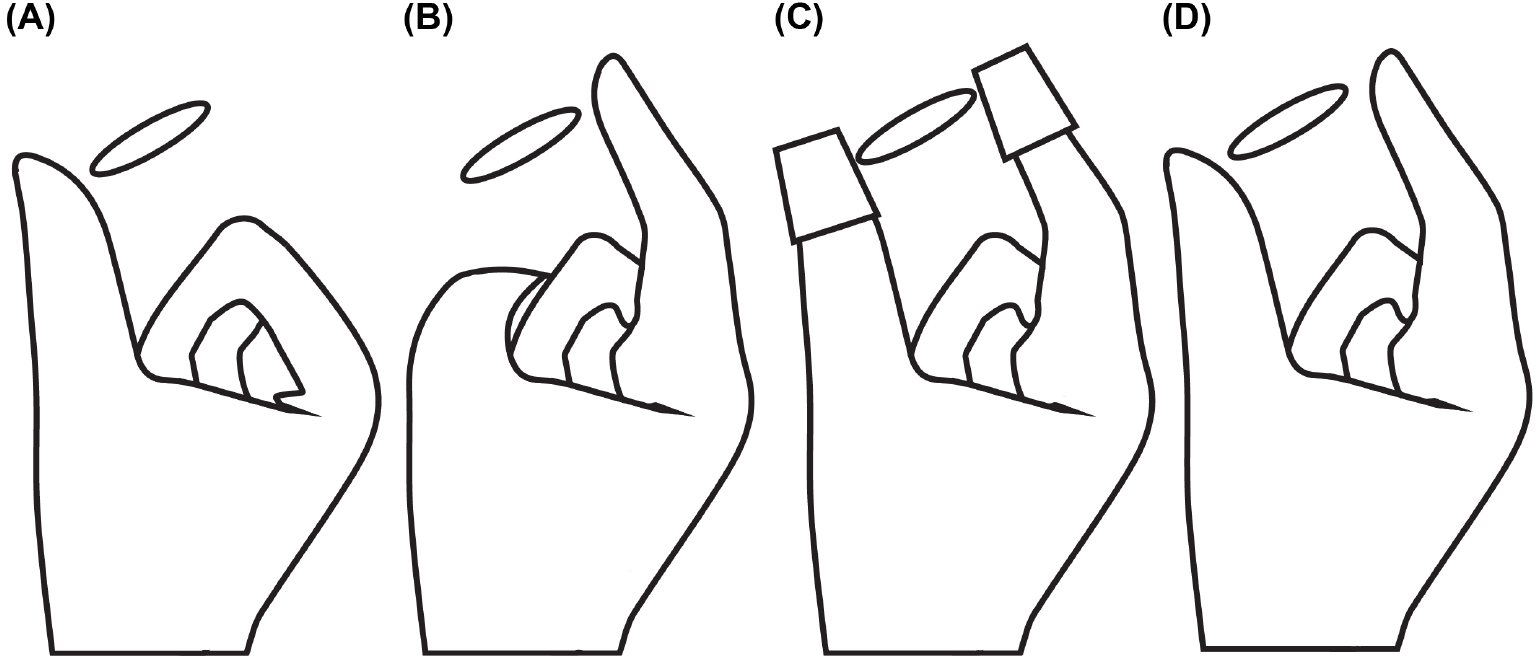
Demonstration of the cue conditions. **(A)** Cut.1 (cutaneous cue on the thumb). **(B)** Cut.2 (cutaneous cue on the index). **(C)** Configuration cue. **(D)** Combined Condition (all cues present).

### 2.4.1 Experiment 1

Experiment 1 took place over two sessions, 2.5 hours each, a week apart. The conditions were partially counterbalanced such that, *Config, Combined*, and *Cut*.*1* or *Cut*.*2* conditions had equal chances of occurring at the beginning, middle, or end of the experiment. Although *Cut*.*1* and *Cut*.*2* conditions always occurred in succession, their order was also randomized. Participants completed two blocks of each condition across two days for a total of four blocks of 40 trials each. Disks in Experiment 1 were circular with the following radii in millimetres: 10, 13, 14, 14.5, 15, 15.5, 16, 17, and 20. The 15mm disk was the reference to be compared with any of the other disks during each trial. In each block, the stimuli were delivered by Method of Constant Stimuli, with each comparison disk being delivered five times in pseudo-random order. Accordingly, each participant was tested for a total of 160 trials, 20 times with each of the 8 comparison disks, in each condition.

### 2.4.2 Experiment 2

Experiment 2 took place in one 2.5-3 hour session. The conditions were partially counterbalanced in the same way as in Experiment 1. Each condition consisted of two blocks of 40 trials. Disks in Experiment 2 were circular with the following radii in millimetres: 5, 8, 9, 9.5, 10, 10.5, 11, 12, and 15. The 10mm disk was the reference to be compared with any of the other disks during each trial. The stimuli were delivered using a Bayesian Adaptive Procedure which allowed us to present the disk on each trial from which the most information about the participant’s psychometric function could be learned [17, 18]. The Bayesian adaptive procedure can be applied to any psychometric function parameterization; we defined our psychometric functions as cumulative normal distributions of the stimulus level, Δ (Eq. 7).

Here, Δ_*m*_ refers to the participant’s internal sensorineural measurement of the size difference between the two disks, which is assumed to result from a random sample drawn from a Gaussian distribution centered on the actual size difference, Δ.

### 2.4.3 Experiment 3

Experiment 3 took place in one 2.5-3 hour session. Comparison disks in Experiment 3 were circular with the following radii in millimetres: 5, 8, 9, 9.5, 10.5, 11, 12, and 15. The methodology used was the same as in Experiment 2 with the following exceptions. Instead of 2 blocks of 40 trials for each of 4 conditions, participants completed one block of 40 trials for each of 7 different conditions. The first 3 conditions were the same as the earlier experiments, to estimate participant sigmas for the individual cues (*Cut*.*1, Cut*.*2*, and *Config*). The remaining 4 conditions are all *Cue Conflict* conditions. The *Cue Conflict* stimuli were designed such that each indicates a curvature and radius in conflict with one another. Figure 5 illustrates the design of **-3** and **+3** conflict stimuli. The magnitude of the number indicates the difference in radius (mm) from the reference, and the sign indicates in which direction the lateral distance across the face of the disk changed. Thus, a **-3** conflict disk (Fig. 5A) has the lateral diameter of a disk with a radius that is −3mm from the reference of 10mm: a 7mm radius. Simultaneously, it has the lateral *curvature* of a disk +3mm from the reference of 10mm: a 13mm radius. A **+3** conflict disk (Fig. 5B) has the lateral diameter of a disk with a radius that is +3mm from the reference of 10mm: a 13mm radius. Simultaneously, it has the lateral *curvature* of a disk −3mm from the reference of 10mm: a 7mm radius. The same logic applies to the **-2** and **+2** conflict disks.

**Fig. 5.**
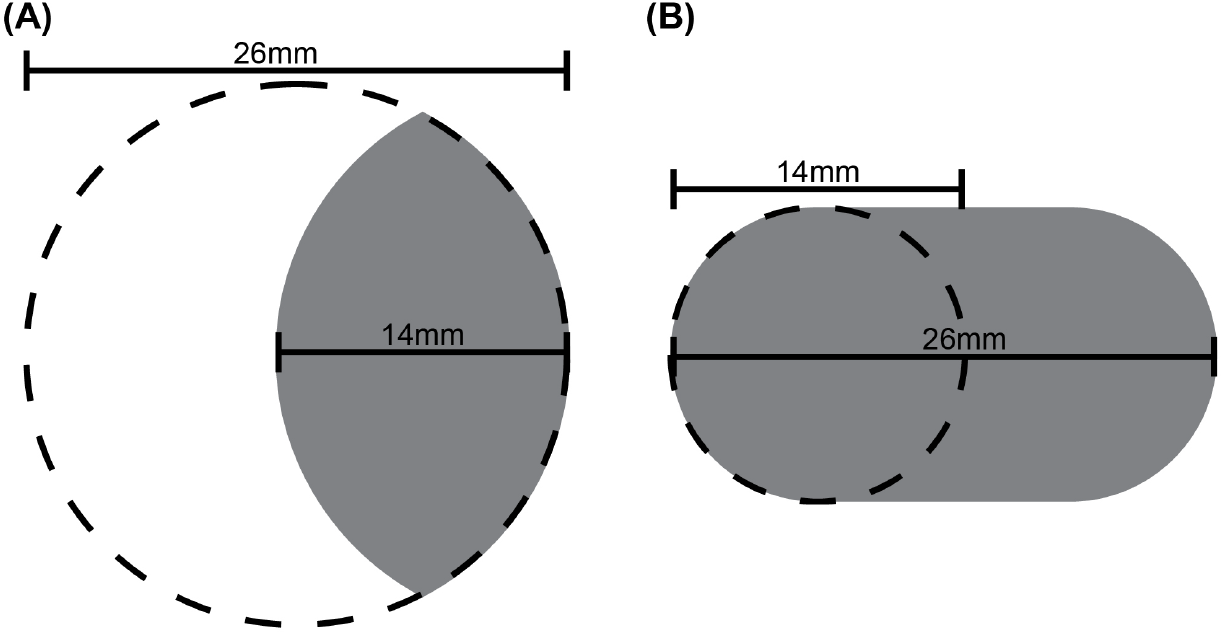
Illustration of two of the four conflict stimuli. Recall that participants felt disks laterally along the horizontal axis. A) The **-3** conflict stimulus (shaded) has a horizontal extent equivalent to that of a **-3 mm** radius disk, and a curvature equivalent to that of a +3 mm radius disk, relative to a 10 mm radius circular disk. B) The **+3** conflict stimulus has a horizontal extent equivalent to that of a **+3 mm** radius disk, and a curvature equivalent to that of a −3 mm radius disk, relative to a 10 mm radius circular disk.

From the participant’s point of view, these were no different from the *Combined* conditions of the earlier experiments. In each of these conflict conditions, however, the 10mm radius reference disk was swapped out for 4 different cue conflict stimuli. The conditions were partially counterbalanced with restrictions: individual cue conditions occurred at the beginning of the experiment, but *Cut*.*1* and *Cut*.*2* conditions occurred successively in random order; conflict conditions occurred at the end of the experiment in random order, but ±**2** conditions occurred together and ±**3** conditions occurred together. The task was otherwise carried out in the same manner as before.

### 2.5 Statistical Analyses

We employed frequentist statistics and Bayesian inferential methods to describe and analyse the data. We discretized sigma into 801 values ranging from 0.01 mm to 40.01 mm and applied a uniform prior over this range. We parameterized each participant’s psychometric function (Equation 7) and estimated sigma as described (Equations 8, 9) For frequentist analyses, the mode of each participant’s posterior PDF over sigma was taken as a point estimate. Each participant’s individual cue sigmas were combined according to each model’s formulas to calculate the model’s prediction for the participant’s combined cue condition. Frequentist statistical analyses were completed using Python and relevant packages and modules including pandas, numpy, statsmodels, and matplotlib.

The Bayesian model comparison between the three hypotheses was calculated in the same manner as for the experiment simulations in 2.1.2 with the marginal likelihood formulas given by equations 10 and 11. From these marginal likelihoods, and using a uniform prior for the models, we calculated the posterior probability of each model for each participant.

## 3 Results

### 3.1 Experiment 1

In Experiment 1, participants compared the sizes of two sequentially presented disks. In different conditions, the participants had access to either a single sensory cue or to the three cues combined. Figure 6 plots the mean single-cue and combined cue sigma estimates across participants, along with the combined cue sigma predicted by each model on the basis of the single cue sigma values. The plot clearly indicates that reliability was higher (i.e., sigma was lower) for the configuration cue than for either of the cutaneous cues. Furthermore, the combined condition had a reliability that was similar to that of the configuration condition.

**Fig. 6.**
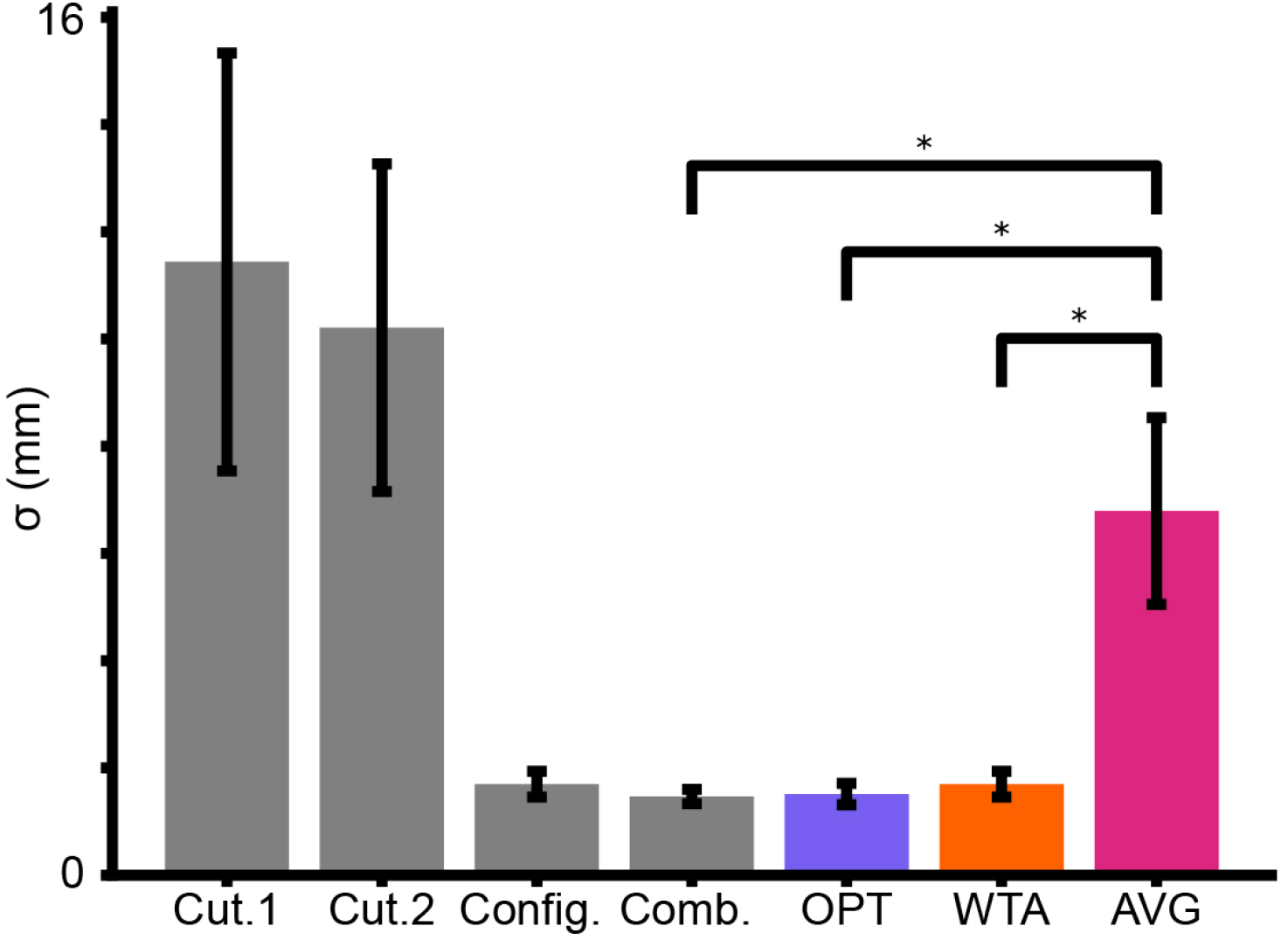
Mean sigma estimates across participants in Experiment 1 (Grey). Mean model predictions (colored bars) for performance in the Combined Condition, based on the Cutaneous and Configuration conditions. Error bars = +/- SE. Asterisk (*) = Significant t-test.

One-way repeated measures ANOVAs reveal significant effects of condition (p = 0.006, F(3, 33) = 4.961) and model prediction (p = 0.001, F(2, 22) = 9.038). Paired-sample t-tests demonstrate that the *AVG* model prediction differs significantly from both the *OPT* (p = 0.010, t(11) = 3.094) and *WTA* model predictions (p = 0.013, t(11) = 2.922). There was, however, no significant difference between the predictions of the *WTA* and *OPT* models (p = 0.050, t(11) = 2.197). Importantly, there was a significant difference between the Combined Condition and the *AVG* model prediction (p = 0.010, t(11) = 3.099) but not between Combined and *OPT* (p = 0.730, t(11) = 0.354) or Combined and *WTA* (p = 0.206, t(11) = 1.343).

Applying Bayesian model comparison, we calculated posterior probabilities for each participant for each model. The individual posterior probabilities for each participant in Experiment 1 are reported in Table 3. Median posterior probabilities corroborate the trends from the above frequentist statistics. The median posterior probabilities are as follows: *OPT* = 0.532, *WTA* = 0.457, *AVG* = 1.39 *·* 10^−3^. Overall, the Bayesian model comparison categorized 8 out of 12 participants as *OPT*, but the posterior probabilities were often quite close for *OPT* and *WTA*.

**Table 3.** Exp. 1 Model Posterior Probabilities by Participant.

| Participant | OPT | WTA | AVG | Classification |
| --- | --- | --- | --- | --- |
| 1 | 0.472 | 0.527 | 0.001 | WTA |
| 2 | 0.458 | 0.540 | 0.002 | WTA |
| 3 | 0.507 | 0.493 | 0.000 | OPT |
| 4 | 0.535 | 0.463 | 0.002 | OPT |
| 5 | 0.548 | 0.447 | 0.004 | OPT |
| 6 | 0.052 | 0.182 | 0.766 | AVG |
| 7 | 0.588 | 0.045 | 0.367 | OPT |
| 8 | 0.481 | 0.519 | 0.000 | WTA |
| 9 | 0.548 | 0.451 | 0.000 | OPT |
| 10 | 0.560 | 0.440 | 0.000 | OPT |
| 11 | 0.665 | 0.118 | 0.217 | OPT |
| 12 | 0.528 | 0.472 | 0.000 | OPT |

### 3.2 Experiment 2

Experiment 1 provided evidence against the *AVG* model but was unable to convincingly distinguish between the *OPT* and *WTA* models. We reasoned that this difficulty may have resulted from the fact that *Config* had a much smaller sigma than *Cut*.*1* and *Cut*.*2* in most participants. In Experiment 2, with the goal of enhancing *Cut*.*1* and *Cut*.*2*, we made the disks smaller so that they would indent the fingers more sharply. We hoped to reduce the sigma of the cutaneous cues so that the cutaneous and configuration cues would have more similar sigma values, facilitating the classification of participants as either *OPT* or *WTA*.

Figure 7 shows a reduction in sigma in the Cutaneous Conditions. The reliabilities of the cues in the Cutaneous and Configuration Conditions are relatively closer together and we begin to observe a greater difference between the predictions of the *OPT* and *WTA* models.

**Fig. 7.**
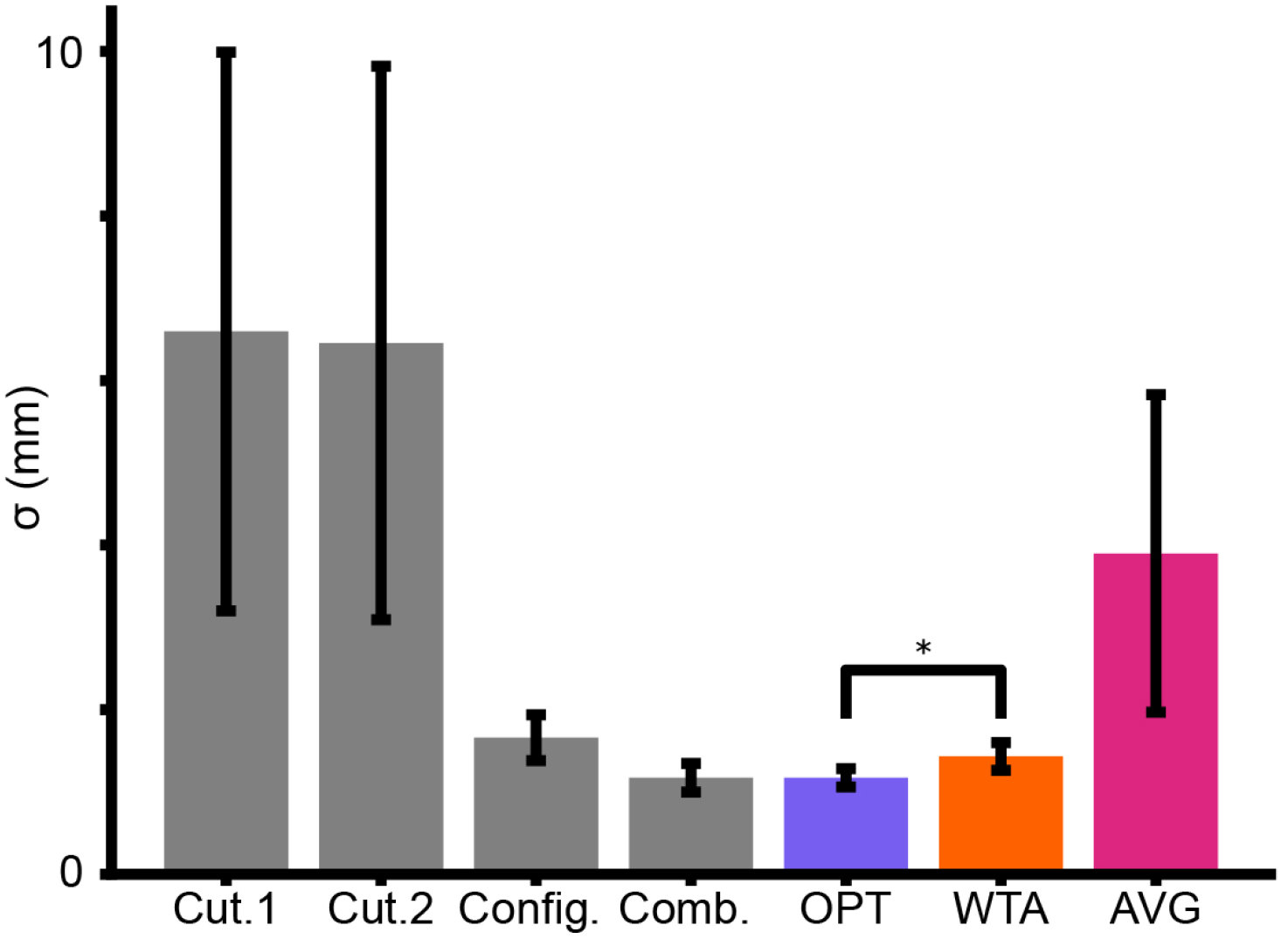
Mean sigma estimates across participants in Experiment 2 (Grey). Mean model predictions (colored bars) for performance in the Combined Condition, based on the Cutaneous and Configuration conditions. Error bars = +/- SE. Asterisk (*) = Significant t-test. Note the difference in y-axis scale from Fig. 6.

One-way repeated measures ANOVAs reveal non-significant effects of condition (p = 0.104, F(3, 30) = 2.236) and model prediction (p = 0.187, F(2, 20) = 1.825).

Calculated posterior probabilities provide additional nuance and context to the above findings. The individual posterior probabilities for each participant in Experiment 2 are reported in Table 4. The median posterior probabilities are as follows: *OPT* = 0.502, *WTA* = 0.393, *AVG* = 0.097). Overall, the Bayesian model comparison categorized 8 of the 11 participants as *OPT*. While the posterior probabilities for OPT and *WTA* were often close, the posterior probabilities from Experiment 2 were generally more strongly in favour of *OPT* over *WTA* than were the posterior probabilities from Experiment 1.

**Table 4.** Exp. 2 Model Posterior Probabilities by Participant.

| Participant | OPT | WTA | AVG | Classification |
| --- | --- | --- | --- | --- |
| 1 | 0.500 | 0.482 | 0.019 | OPT |
| 2 | 0.188 | 0.385 | 0.427 | AVG |
| 3 | 0.010 | 0.659 | 0.331 | WTA |
| 4 | 0.199 | 0.370 | 0.430 | AVG |
| 5 | 0.838 | 0.131 | 0.031 | OPT |
| 6 | 0.455 | 0.437 | 0.109 | OPT |
| 7 | 0.773 | 0.044 | 0.183 | OPT |
| 8 | 0.538 | 0.462 | 0.000 | OPT |
| 9 | 0.720 | 0.269 | 0.011 | OPT |
| 10 | 0.502 | 0.498 | 0.000 | OPT |
| 11 | 0.524 | 0.362 | 0.114 | OPT |

### 3.3 Experiment 3

For Experiment 3, we incorporated a cue-conflict paradigm. Figure 8 shows an overview of participant performance in each experimental condition. Observing the mean sigma estimates across participants for each condition, we found similarities to earlier experiments. The mean sigma of the cues decreased from *Cut*.*1* to *Cut*.*2* to *Config* The four Cue Conflict Conditions are analagous to the earlier Combined Conditions of previous experiments because participants have access to all cues. As an aside, sigmas in the Conflict Conditions generally appear to be lower than the the sigmas of any individual cue, with the exception of the particularly noisy **+3** Condition. Nevertheless, Experiment 3 was not intended as a replication. The predictions and planned analyses were different than in the earlier experiments.

**Fig. 8.**
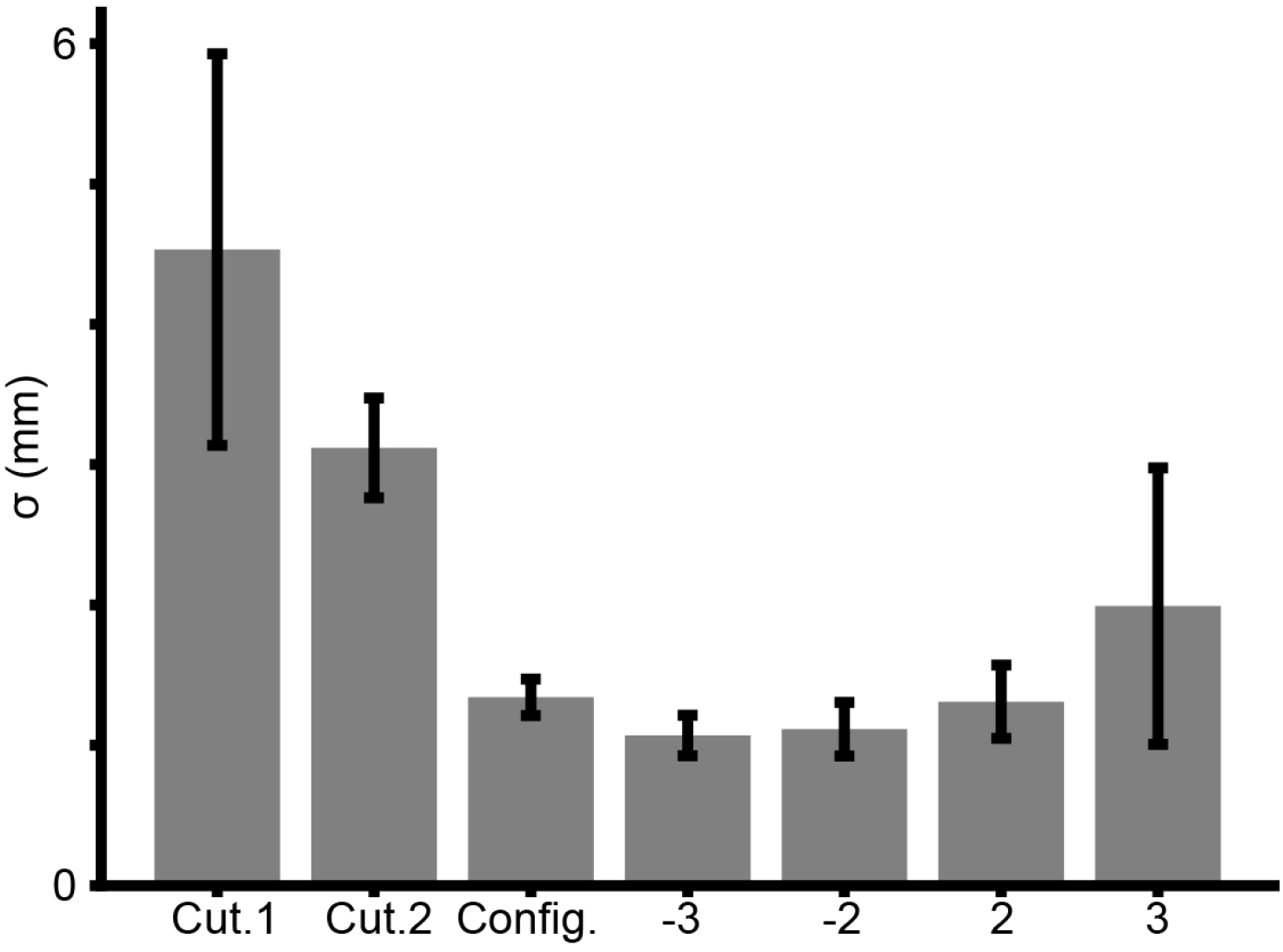
Mean sigma estimates across participants in Experiment 3 for the individual cue conditions and conflict conditions. Error bars = +/- SE.

As the reference in each of the Conflict Conditions is one of four conflict disks, each model predicts that the *PSE* should shift linearly in the direction of the cues upon which the participant most relies. For example, if a participant is relying solely on *Config*, a **+3** conflict disk would feel subjectively equal to a 13mm radius disk.

Since, in a *2IFC* task, the response is to which disk felt larger, the participant would be guessing when comparing the **+3** conflict disk to a 13mm comparison disk. On average, the proportion of times one disk is reported to be greater than the other would be 0.5: a rightward shift of 3mm in the *PSE*.

The mean *PSE* shift across participants, as a function of conflict condition, is shown in Figure 9. Each model predicted a different linear *PSE* shift. The *AVG* model predicted a negative slope because of the influence of both *Cut*.*1* and *Cut*.*2*, which produced measurements counter to *Config*. The *WTA* model predicted a positive slope equal to 1 for any participant for whom *Config* was best and the *PSE* shift would be equal to the Conflict Condition. On average, the shifting *PSEs* most closely match the predictions of the *OPT* model. The prediction of the slope for the optimal model was based on Equation 12 where *s* is the Conflict Condition and *ŝ* is the PSE. The positive slope indicates that *Config* was more heavily relied upon than *Cut*.*1* or *Cut*.*2*. Importantly, the slope is not equal to 1, which implies that *Cut*.*1* and *Cut*.*2* did indeed influence perception and cues were combined.

**Fig. 9.**
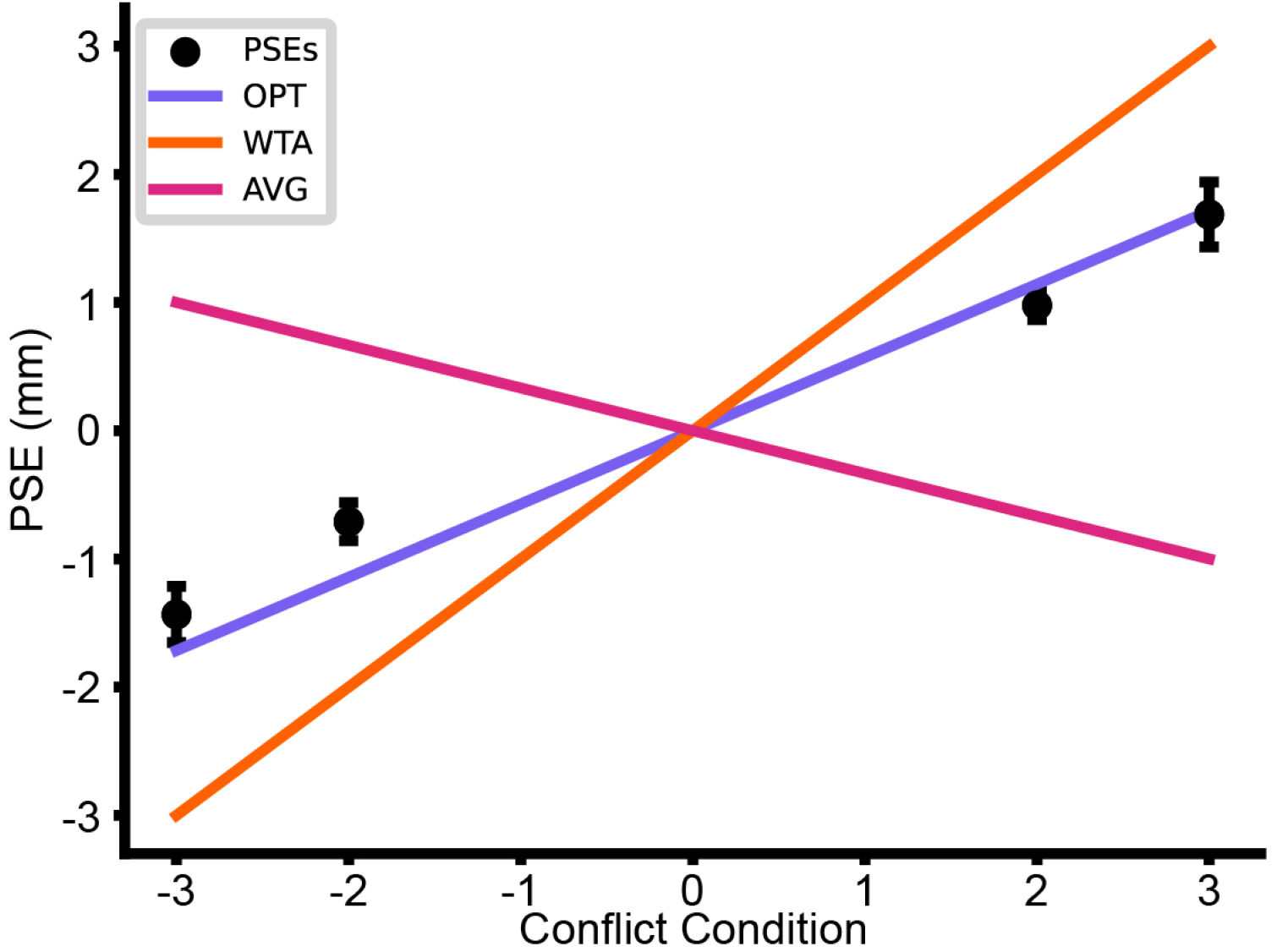
Mean PSE shift across participants by Conflict Condition (Error bars = *±* SE)

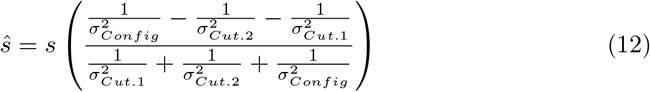

The *PSE* shifts, however, are still likely to be positive if *Config* was truly more relied upon. Notably, the actual recorded PSEs of each participant have positive slopes even if the model prediction was negative. This matched our results from Figure 8 and earlier experiments, where *Config* has the lowest sigma of any individual cue, on average. Thus, participants appear to combine these cues, relying on *Config* more than the others, although *Cut*.*1* and *Cut*.*2* generally have influence on perception as well. Table 5 displays the posterior probabilities for each participant across the three models, rounded to two decimal places. The majority of participants were strongly *OPT* over *WTA* or *AVG*. Participant 11, however, appeared to be strongly *WTA*.

**Table 5.** Exp. 3 Model Posterior Probabilities by Participant.

| Participant | OPT | WTA | AVG | Classification |
| --- | --- | --- | --- | --- |
| 1 | 1.000 | 0.000 | 0.000 | OPT |
| 2 | 1.000 | 0.000 | 0.000 | OPT |
| 3 | 1.000 | 0.000 | 0.000 | OPT |
| 4 | 0.999 | 0.001 | 0.000 | OPT |
| 5 | 1.000 | 0.000 | 0.000 | OPT |
| 6 | 0.994 | 0.006 | 0.000 | OPT |
| 7 | 0.999 | 0.000 | 0.001 | OPT |
| 8 | 0.973 | 0.027 | 0.000 | OPT |
| 9 | 0.871 | 0.001 | 0.127 | OPT |
| 10 | 1.000 | 0.000 | 0.000 | OPT |
| 11 | 0.000 | 1.000 | 0.000 | WTA |

## 4 Discussion

The present study reported three experiments that contribute increasingly supportive evidence that humans optimally combine hand configuration and cutaneous cues to perceive the size of a disk grasped between the index finger and thumb. Experiment 1 provided insight into the relative reliability of the three cues. The data suggested an

*Exceptional Sigma* scenario in which *Config* was much better than *Cut*.*1* and *Cut*.*2*. In such a scenario, as shown in our simulations, the *WTA* and *OPT* models make similar predictions. Accordingly, while the data from Experiment 1 provided strong evidence against the *AVG* model, we were unable to convincingly differentiate between the *WTA* and *OPT* models. In Experiment 2, we aimed to increase the reliability (lower the sigmas) of the cutaneous cues to better differentiate between the *WTA* and *OPT* models. we hypothesized that the use of smaller disks would result in greater skin deformation and correspondingly more reliable cutaneous cues [16] while leaving the configuration cue relatively unaffected. Our intention was that this would cause *OPT*, but not *WTA*, to predict a lower combined condition sigma, permitting us to differentiate between these two models. Experiment 2 demonstrated that reducing disk size indeed preferentially reduced the sigmas of the cutaneous cues. Our inferences 13 also became more in favour of *OPT* over *WTA*. In Experiment 3, we concluded our investigation with a cue conflict experiment. By using stimuli that put the cutaneous cues in conflict with the configuration cue, we were able to glean information from *PSE* shifts. The *PSEs* showed that participants generally rely more on *Config* than *Cut*.*1* or *Cut*.*2* but information from all cues is integrated in a way consistent with *OPT*. Our inferences in Experiment 3 were notably confident in their classifications. These results mirror the results of our model simulations, lending credence to the conclusions. Taken together, these experiments support the conclusion that humans optimally integrate cutaneous and proprioceptive cues for haptic size perception.

### 4.1 Comparison to Previous Literature

Our findings are generally consistent with previous multimodal studies and unimodal studies outside of haptics. Ernst and Banks [5] found that participants could optimally combine visual and haptic cues for height. Gepshtein and Banks [9] found participants were close to statistically optimal in a visuo-haptic distance discrimination task. Alais and Burr [3] demonstrated optimal cue combination across auditory and visual stimuli. When localizing a spatial source by a beep and a flash, participants were better with both cues than with either alone. On some trials, they introduced cue conflicts where the beep and flash were displaced in opposite directions. Participants judged the location of the event to be closer to whichever of the two cues was more reliable. Hillis et al. [4] demonstrated optimal cue combination for two visual cues to depth: a texture cue created with Voronoi patterns viewed monocularly, and a disparity cue using random dots viewed binocularly. Negen et al. [11] found participants, with training, could integrate a newly learned sensory skill with vision in a statistically optimal manner. In virtual reality spatial navigation and memory tasks, Newman and McNamara [19] found that participants could combine visual landmark cues optimally, depending on task complexity and perspective. Many studies in these domains find that the reliability of cues is considered during the process of perception [4–11].

In some experimental paradigms, in non-tactile modalities such as vision, stimulus reliability can be adjusted during an experimental session. Given the need for physical stimulus delivery, this can be challenging in a tactile experiment. Through the use of virtual haptic or force-feedback devices, however, it is possible to adjust certain aspects of a haptic stimulus, such as the distance between fingers. Simultaneously adjusting a meaningful cutaneous cue, however, would be substantially more challenging. Instead, our study made use of 3D-printed disks and a calibrated mechanical system on which to mount them. As such, we carefully selected our stimulus range to fit the limited space available. We correctly hypothesized that smaller disks would make for better cutaneous cues while leaving the hand configuration cue relatively unaffected. Logistical and engineering considerations, however, prevented us from manipulating stimulus reliability throughout the course of a single experiment.

Overall, our haptic cue combination results share the general trends of the broader literature. The consistency our results show with previous studies suggests that the brain integrates multiple tactile cues in a similar fashion as it does with other sensory modalities.

### 4.2 Future Directions

Our experiments revealed some interesting examples of individual variability. For example, in Experiment 3, Participant 11 was more strongly *WTA* than *OPT* or *AVG*. Future experiments could examine the degree or source of such individual variability.

An interesting question for future investigation is whether a participant who begins strongly WTA could shift towards OPT with sufficient training.

We reduced disk size to lower the cutaneous sigmas and bring their performance closer to *Config* performance. A future study could aim to lessen the disparity between cutaneous and configuration cues by preferentially worsening *Config* and making its sigma higher. Potentially, this could be accomplished by numbing or vibrating select areas of the hand, but this manipulation would not be trivial. Ideally, if the manipulations resulted in each of the three cues having *Similar Sigmas*, then the difference between the predictions of the *WTA* and *OPT* models would be at their greatest.

Future experiments could also investigate heteroscedasticity in the sensory signal.

As a first approximation, our analyses assumed that participants have a single value for sigma for each cue. It is possible, however, that the trend we observed of increasing sigma, as conflict increases (Figure 8), reflects a Weber fraction [20]. This possibility could be investigated with experiments that use a much larger stimulus size range.

In our experiments, from one interval to another within a trial, there was a short delay between stimulus presentations. Thus, participants had to engage working memory to hold in mind the perceived size of the first disk in order to compare it to the second. Would participant performance improve, or perhaps worsen, if we were to deliver the reference to one hand and the comparison to the other, simultaneously?

Our experiments involved controlled haptics. The hand moved towards and grasped the target, but participants could not freely explore the stimuli. Future experiments could reduce the level of control in favor of a more naturalistic setting. If participants were provided unfettered access to the stimulus objects and allowed to grasp and scan the disks with all of their fingers and perhaps both of their hands, how much might performance improve? Would the *Config* cue still be most informative?

Better understanding the means by which humans combine multiple haptic cues will ultimately support advancements in fields such as haptic VR, robotics, and telesurgery. Our research points towards the relative importance of proprioceptive and cutaneous cues. While cutaneous cues were less reliable overall, they measurably improved performance even when hand configuration was more reliable. This highlights the importance of utilizing haptic (including cutaneous) feedback in areas like telesurgery, where incremental gains in performance can lead to invaluable improvements in outcomes. Additionally, fields such as haptic VR are constrained by the cost, size, and resolution of tactile stimulators. Cue combination research may suggest means of cleverly employing multiple cues to effectively enhance perceptual resolution within the constraints of available hardware.

## Author Contributions

Keon S. Allen: Writing - original draft, Writing - review and editing, Conceptualization, Investigation, Data curation, Methodology, Formal analysis, Visualization.

Daniel Goldreich: Writing - original draft, Writing - review and editing, Conceptualization, Investigation, Data curation, Methodology, Project administration, Formal analysis, Supervision, Funding acquisition, Visualization.

## Acknowledgments

We thank Kyra Gauder for helpful discussion and technical assistance, Bradley Karat and Maraam Haque for their assistance with data collection, and Khesraw Mohibi and Arjun Patel for discussion regarding simulations.

